# Differentiated kidney tubular cell-derived extracellular vesicles enhance maturation of tubuloids

**DOI:** 10.1101/2022.02.08.479621

**Authors:** Rafael Soares Lindoso, Fjodor A Yousef Yengej, Franziska Voellmy, Maarten Altelaar, Estela Mancheño Juncosa, Theano Tsikari, Carola M.E. Ammerlaan, Bas W.M. Van Balkom, Maarten B Rookmaaker, Marianne C Verhaar, Rosalinde Masereeuw

## Abstract

Advanced kidney *in vitro* models such as organoids or tubuloids still lack the intrinsic expression of various transport proteins needed for active secretory function. Extracellular vesicles (EVs), cell-derived structures that constitute the organ’s microenvironment, are known to regulate various cellular processes, including kidney development and regeneration across the nephron. In this study, we propose a new application of renal tubular epithelial cell EVs as modulators for tubuloid functional maturation by increasing the levels of various differentiation markers such as organic anion transport 1 (OAT1), a protein involved in endogenous waste excretion. First, we show that EVs from engineered proximal tubule cells increased the expression of several transcription factors and epithelial transporters in tubuloids that resulted in improved cellular transport capacity. Next, a more in-depth proteomic data analysis demonstrated that EVs can trigger various biological pathways, including mesenchymal-to-epithelial transition, which is crucial in the tubular epithelial maturation process. Moreover, we demonstrated that EV-treated tubuloid-derived cells in a 3D tubular conformation as part of a bioartificial kidney can generate a tight polarized epithelial monolayer with formation of dense cilia structures. In conclusion, EVs from renal tubular epithelial cells can phenotypically improve tubuloid maturation, thereby enhancing their potential as preclinical models and functional units in regenerative therapies.

## 1. INTRODUCTION

End-stage kidney disease (ESKD) is among the main causes of morbidity and mortality worldwide [1]. When kidney function declines below 10-15%, solutes accumulate in blood and cause toxicity, affecting the functional capacity of the kidney and other organs in the body [2]. Although kidney transplantation is presented as best solution for ESKD patients, the organ viability does not cope with the demand. In addition, the contribution of dialysis treatment, cannot fully substitute normal kidney function and bring several drawbacks for the patients in long-term chronic dialysis [3]. In order to improve patient’s life, new strategies have been developed to use bioartificial kidney as replacement therapy [4,5]. The use of cellular components within the bioartificial kidney has brought the possibility to mimic different kidney aspects like transport, metabolic and endocrine functions [6,7]. However, to translate such advances into clinic, is critical to define a cell source that can represent the different specialized cell types present in the kidney and, at the same time, be biocompatible with the patient.

Organoids are three-dimensional structures derived from stem cells that reproduce *in vitro* the formation of near-physiological tissues [8]. Due to their self-renewal and self-organizational capabilities, organoids are state of the art cell culture models to study organ development and disease, especially for organs with higher complexity, such as the kidney. After differentiation of (induced) pluripotent stem cells towards the two kidney precursor populations (i.e., ureteric bud and metanephric mesenchyme), reciprocal interaction causes self-organization and patterning to generate nephron structures [9]. An interesting alternative cell source to pluripotent stem cells are kidney progenitor cells derived from adult kidneys; upon culturing, these tissue models, defined as tubuloids, do not form nephron structures but instead allow for long-term expansion and generation of primary functional renal epithelium reflecting several aspects of all tubular regions of the mature nephron [10,11]. Tubuloids are a promising tool for kidney disease modeling and personalized drug screening, but also for regenerative medicine purposes, as adult progenitors can be patient-derived and do not require any genetic modification [10–12]. However, despite being more mature than (induced) pluripotent stem cell-derived organoids, tubuloids do not yet present a fully developed phenotype, especially for proximal tubules [10]. This is a common shortfall observed in kidney cells cultured *in vitro*: thus far, no existing proximal tubule cell model has shown a full intrinsic physiological recapitulation of the native tissue, including transepithelial transport capacity, limiting their application in pharmacological and toxicological studies [13,14].

The ability of the kidney to actively excrete metabolic waste, drugs and their metabolites is given by the presence of transporters in the membranes of proximal tubule cells [12]. Among these, organic anion transporter 1 (OAT1) is highly expressed that together with apical efflux pumps like multidrug resistance-associated proteins (MRPs) and breast cancer resistance protein (BCRP) contributes to the transfer of a large range of organic compounds from the blood circulation to the urine, including drugs and endogenous metabolites, in a highly controlled manner [15,16]. As OAT1 is an important determinant of drug induced kidney injury [17,18], its absence limits the application of kidney-derived cells, organoids and tubuloids as drug screening platforms in the early phases of drug development but hampers also other applications such as in regenerative therapies. Thus, improving kidney cell maturation *in vitro* to reach expression of OAT1 and other proximal tubular transporters to levels close to native tissue is crucial for their predictive capacity as a preclinical test platform [19].

Strategies to improve kidney organoid and tubuloid maturation have focused on the use of different growth factors and aspects of the kidney microenvironment (e.g., 3D organization, vascularization, extracellular matrix, or fluid flow) [20–23]. In this context, the use of extracellular vesicles (EVs) has been described as a suitable strategy in regulating kidney regeneration [24,25]. These vesicles are nanosized lipid bilayer structures that mediate cellular communication through the transfer of bioactive molecules, including proteins, nucleic acids and lipids [26]. Based on biogenesis, size and content, EVs can be classified as exosomes (30 — 150 nm), microvesicles (100 nm — 1 μm) and apoptotic bodies (50 nm — 5 μm), presenting different biological functions based on their cargo [27]. Kidney EVs participate in intranephron communication *in vivo* by transferring unique proteins and RNAs that can modulate the functional activity of tubular epithelial cells [18,19]. For instance, urinary EVs isolated from rats and from conditioned medium (CM) of proximal tubule cells contain functionally active aquaporin 2 (AQP2), which can be transferred to other tubular epithelial cells [24,25]. EVs secreted by human kidney proximal tubule cells were also shown to modulate the regulation of purinergic signaling in collecting duct cells through increased extracellular ATP and downregulation of epithelial sodium channel (ENaC) [26]. Moreover, CM and isolated EVs from kidney epithelial cells were capable of inducing mesenchymal cells to acquire a more epithelial phenotype [28–30]. Together, these data clearly show that EVs from renal tubular epithelial cells mediate a complex regulatory communication system that holds the potential to functionally influence tubular epithelial cells and their transport capacity.

As EVs can transfer the imprinting of originator cells to recipient cells, we investigated the potential of EVs derived from differentiated kidney proximal tubule cells overexpressing OAT1 to support tubuloid functional maturity by inducing OAT1 expression. First, we describe increased expression of important proximal tubule transcription factors and transport proteins. Next, EV’s cargo was evaluated on proteome basis to elucidate the underlying molecular mechanisms involved in the differentiation process. Finally, the capacity of EVs to functionally mature tubuloids-based bioengineered tubules as functional units of a bioartificial kidney device was investigated.

## 2. MATERIAL AND METHODS

### 2.1 Cell and tubuloids culture

The tubuloids, derived from human cortical kidney tissue, were obtained from Hubrecht Organoid Technology (HUB), Utrecht, the Netherlands (OSR-2020-30b), and were cultured according to Schutgens et al. [10]. Briefly, the tubuloids were maintained at 37°C and 5% v/v CO2 in Basement Membrane Extract (BME) (R&D Systems, Abingdon, UK) and cultured in expansion medium (ADMEM/F12 supplemented with 1% penicillin/streptomycin, HEPES, GlutaMAX, N-acetylcysteine (1 mM; Sigma-Aldrich, Zwijndrecht, the Netherlands) and 1.5% B27 supplement (Gibco, Life Technologies, Paisley, UK), supplemented with 1% Rspo3-conditioned medium (U-Protein Express, Utrecht, The Netherlands), EGF (50 ng ml^−1^; Peprotech, London, UK), FGF-10 (100 ng ml^−1^, Peprotech, London, UK), Rho-kinase inhibitor Y-27632 (10 μM; Abmole, Brussels, Belgium) and A8301 (5 μM; Tocris Bioscience, Abingdon, UK)). For tubuloids differentiation, the medium was changed to ADMEM/F12 supplemented with 1% penicillin/streptomycin, HEPES and GlutaMAX, defined as differentiation medium and the tubuloids were maintained in culture for 7 days.

Human conditionally immortalized proximal tubule epithelial cells that constitutively express organic anion transporter 1 (ciPTEC-OAT1; Cell4Pharma, Oss, The Netherlands) were maintained at 33°C and 5% v/v CO_2_ to proliferate, until reaching up to 90% confluency, in Dulbecco’s modified eagle medium/HAM’s F12 (Gibco), supplemented with 5 μg/ml insulin, 5 μg/ml transferrin, 5 μg/ml selenium, 35 ng/ml hydrocortisone, 10 ng/ml epidermal growth factor, 40 pg/ml tri-iodothyronine (Merck/Millipore Watford, Hertfordshire, UK), and 10% fetal calf serum (FCS; Greiner Bio-One, Alphen aan den Rijn, the Netherlands) [18,31]. Posteriorly, the cell culture was maintained for 7 days at 37°C and 5% v/v CO_2_, using the same medium composition to allow differentiation, expression of OAT1 and monolayer formation, referred to as maturation.

### 2.2 Conditioned medium (CM) production and EVs isolation

After maturation of ciPTEC-OAT1, the cell culture was washed 3 times with Hanks’ Balanced Salt solution (HBSS; Gibco, Life Technologies, Paisley, UK) and maintained for 24 h at 37°C and 5% v/v CO_2_ in tubuloid differentiation medium. The supernatant was collected, centrifuged at 3,000 × *g* at 4°C and filtered under sterile conditions using a 0.22 μm filter (Millex-GV, PVDF; Merck/Millipore Watford, Hertfordshire, UK), and is referred to as CM-OAT1.

To isolate the EVs from ciPTEC-OAT1, the CM-OAT1 was ultrafiltered using Amicon® Ultra-15 Centrifugal Filter Unit with 100 KDa cutoff (Merck/Millipore Watford, Hertfordshire, UK). The final sample with the concentrated EVs, defined as EV-OAT1, was then stored at −80 °C. The size and number of EV-OAT1 were assessed by Nanoparticle tracking analysis using NanoSight NS500 (Malvern, Worcestershire, UK) with the following settings: camera CMOS, laser Blue 405, camera level 13 (NTA 3.0 Levels), slider shutter 800, FPS 25, number of Frames 1500 and temperature 24.5 - 24.6 °C. To determine the specificity of EV-OAT1 actions, the concentrated EVs were further isolated from the remaining medium using ExoQuick-TC (SBI System Bioscience, Palo Alto, CA, USA). The isolation was performed accordingly to the manufacturer’s protocol, maintaining a proportion of 5 ml of EV’s concentrated medium to 1 ml of ExoQuick-TC and incubating for 16 h at 4°C. After incubation, the samples were centrifuged at 1,500 × *g* for 30 min at 4°C, the supernatant was removed, and a second round of centrifugation at 1,500 × *g* for 5 min at 4°C was performed to remove the remaining supernatant. The isolated EV-OAT1 were used for incubation with tubuloids with a final concentration equivalent to the CM, performing the analysis of RNA content, Western blotting (WB) and further characterization by proteomic analysis.

### 2.3 Tubuloid differentiation with CM or EVs

To investigate the role of CM-OAT1 and EV-OAT1, tubuloids initially cultured in 12-well plates in the presence of expansion medium were divided into 3 experimental groups (Figure 1a): i) control condition (TUB CTR), tubuloids cultivated for 7 days in the presence of differentiation medium, changed every 48 h (standard protocol); ii) tubuloids cultured with CM-OAT1 (TUB CM-OAT1) for 7 days with CM-OAT1 changed every 48 h (3 changes in total); iii) tubuloids cultured with EV-OAT1 (TUB EV-OAT1) for 7 days in the presence of EVs derived from ciPTEC-OAT1 (5×10^8^ particles/well each stimulation), with every 48 h a new stimulation with EVs (3 stimuli in total). In all experimental conditions, the tubuloids were cultured at 37°C and 5% v/v CO_2_. After the 7 days of differentiation, tubuloid cultures were washed 3 times with cold HBSS and transferred to a 15 ml tube in cold HBSS and centrifuged (500 × *g* at 4°C, 5 min). The supernatant containing HBSS and the excess of BME was removed. The pellet containing tubuloids were then used for further analysis.

**Figure 1.**
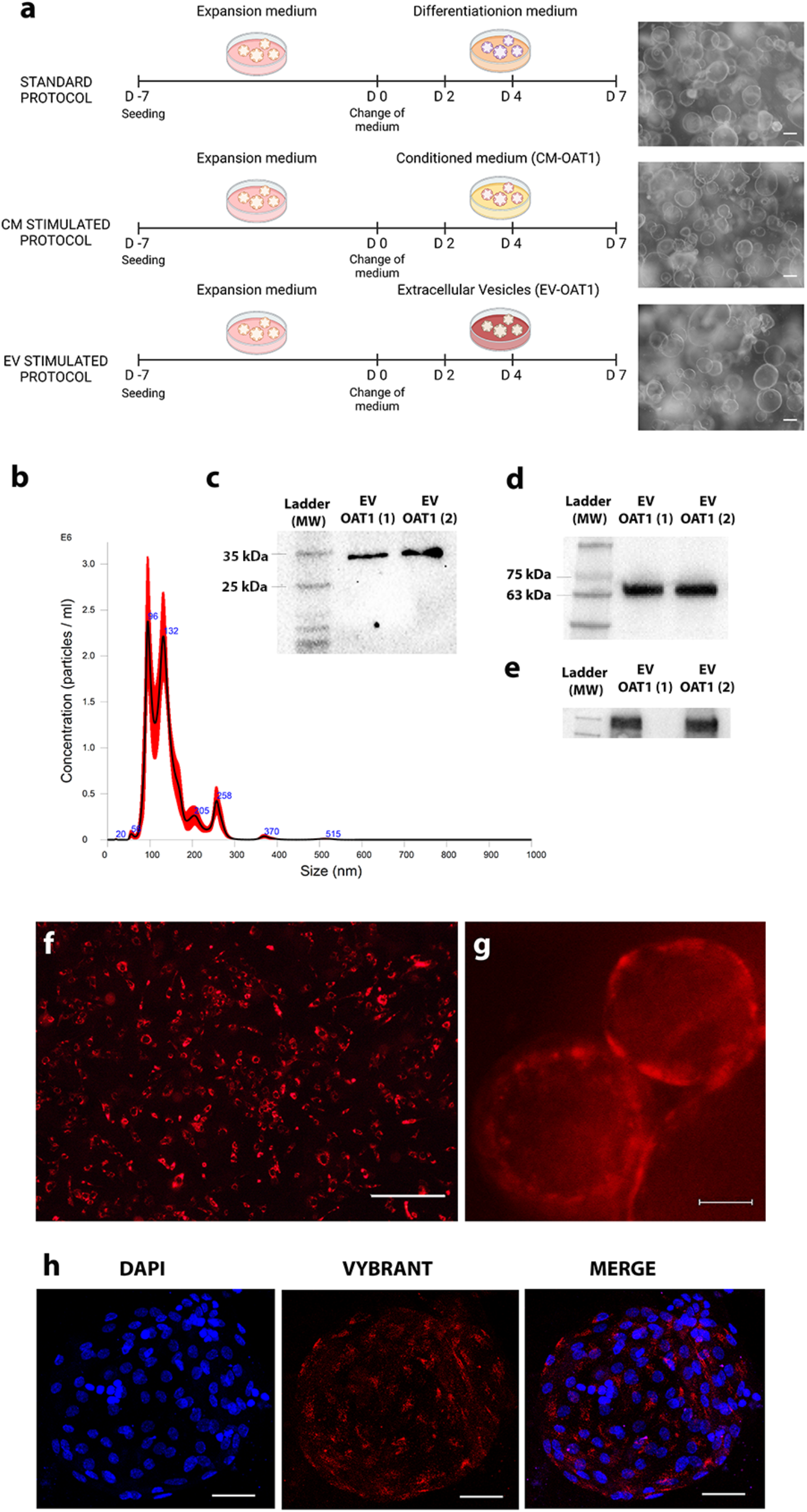
Characterization of EV-OAT1 and uptake by tubuloids. (a) Scheme of the tubuloid differentiation protocols used. The days (D) between D-7 to D0 comprehend the expansion phase of tubuloids. D0 -D7 regards the expansion phase. D0, D2 and D4 indicate the days where new stimulation was given (with differentiation medium, CM-OAT1 or EV-OAT1). (b) Nanoparticle Tracking Analysis representative graph of EV-OAT1. The graph shows the size distribution of EVs (abscissa) and their concentration in particles/ml (ordinate). (c) Representative Western blotting showing the presence of CD63 as exosome marker in EV-OAT1. (d) Representative Western blotting showing the presence of OAT1 within EV-OAT1 cargo. (e) Representative Western blotting showing the presence of Na^+^/K^+^-ATPase within EV-OAT1 cargo. (f) Fluorescence image of fully differentiated ciPTEC-OAT1 culture stained with Vybrant DiI (in red) (scale bar = 500 μm). (g) Fluorescence image showing the uptake of stained EV-OTA1 (in red) by tubuloids after 24 h incubation (scale bar = 50 μm). (h) Representative confocal image of EV-OAT1 distribution into tubuloids after 24 h incubation. The nuclei of the cells were stained with DAPI (in blue) and the EVs were stained with Vybrant DiI (in red) (scale bar = 50 μm).

### 2.4 EVs uptake by tubuloids

To evaluate the EVs incorporation by tubuloids, matured ciPTEC-OAT1 were labelled with Vybrant® DiI (Thermo Fisher Scientific, Vilnius, Lithuania) and EVs, also labelled, were isolated from the supernatant of these cells [32,33]. The tubuloids were incubated for 24 h with the labelled EVs. After, the tubuloids still inside the BME were washed 3 times with cold HBSS followed by a 20 min incubation with Dispase solution (StemCell Technologies, Cambridge, UK) at 37°C and 5% v/v CO_2_ to remove the BME around the tubuloids. The tubuloids were then fixed with 4% paraformaldehyde. Nuclei were stained with ProLong™ Gold antifade reagent containing DAPI (Life Technologies, Eugene, OR, USA). The kidney EV incorporation was analyzed by inverted fluorescence microscopy (Nikon Eclipse Ts2, Melville, NY) for fresh cultures. For fixed cultures, images were acquired using confocal microscopy (Leica TCS SP8 X, Leica Biosystems, Amsterdam, The Netherlands) using the software Leica Application.

### 2.5 Protein purification and mass spectrometry by RP-NanoLC–MS/MS

Protein purification and digestion for proteomic analysis were initially given by lysing the samples in a detergent-based buffer – 1% sodium deoxycholate (SDC), 10 mM Tris (2-carboxyethyl) phosphine (TCEP), 10 mM Tris, 40 mM chloroacetamide) – with Complete mini EDTA-free protease inhibitor cocktail (Roche, Woerden, the Netherlands). The samples were boiled for 5 min at 95°C, 50 mM ammonium bicarbonate was added, and digestion was allowed to proceed overnight at 37°C using trypsin (Promega, Madison, WI, USA) and LysC (Wako, Richmond, VA, USA) at 1:50 and 1:75 enzyme:substrate ratios, respectively. The digestion was quenched with 10% formic acid and the resulting peptides were purified using the Oasis PRiME HLB system (Waters, Wilmslow, UK). Peptides were dried entirely and then resolubilised in a 2% formic acid (FA) MS loading buffer. The proteomic analysis was performed by Reversed-Phase Liquid Chromatography Mass Spectrometry (RP-NanoLC–MS/MS). Data were acquired using an Ultimate3000 system (Thermo Fisher Scientific, Vantaa, Finland) coupled to an Orbitrap Q Exactive HF-X mass spectrometer (Thermo Fisher Scientific, Vantaa, Finland). Peptides were first trapped (Acclaim PepMap100 C18, 5 μm, 100A; Thermo Fisher Scientific, MA, USA) before being separated on an analytical column (Agilent Poroshell, CA, USA; EC-C18, 2.7 μm, 50 cm × 75 μm; Agilent). Trapping was performed for 2 min in solvent A (0.1 M FA in water), and the gradient was as follows: 9–13% solvent B (0.1 M FA in 80% ACN) in 3 min, 13-44% in 95 min, 44–95% in 3 min, and finally 100% for 4 min. The mass spectrometer was operated in data-dependent mode. Full-scan MS spectra from m/z 375–1,600 were acquired at a resolution of 60,000 at m/z 400 after accumulation to a target value of 3 × 10^6^. Up to 15 most intense precursor ions were selected for fragmentation. HCD fragmentation was performed at a normalized collision energy of 27 after accumulation to a target value of 1 × 10^5^. MS/MS was acquired at a resolution of 30,000.

For data analysis, raw mass spectrometry data files were searched using MaxQuant v. 1.6.17.0, against the human Uniprot protein database using Andromeda as a search engine [34]. Cysteine carbamidomethylation was set as a fixed modification and methionine oxidation, protein N-term acetylation. Trypsin was specified as enzyme and up to two miss cleavages were allowed. Filtering was done at 1% false discovery rate (FDR) at the protein and peptide level. Label-free quantification (LFQ) was performed and quantified data were processed and analysed using R and Perseus [35]. The mass spectrometry proteomics data have been deposited to the ProteomeXchange Consortium via the PRIDE [36] partner repository with the dataset identifier PXD027142”. Enrichment analysis related to the identified proteins were performed with FunRich software [37].

### 2.6 Tubuloids functional assay by fluorescein uptake

Evaluation of tubuloid functional transport capacity mediated by OAT1 was evaluated by the uptake of the substrate fluorescein [14]. Initially, tubuloids were removed from the surrounding BME by incubation with Dispase solution (20 min, at 37°C and 5% v/v CO_2_). The tubuloids were then dissociated in single cells by incubation with Accutase® solution (Invitrogen, Carlsbad, CA, USA) for 10 to 15 min at 37°C and 5% v/v CO_2_. The single cells derived from tubuloids were seeded in 6-well plates and maintained in expansion medium until 90% confluency. After the differentiation step of the respective experimental conditions (Figure 1a), the cell cultures were incubated with 1 μM fluorescein (Sigma-Aldrich, MO, USA) for 10 min at 37°C and 5% v/v CO_2_., in the presence or absence of 100 μM Probenecid (Sigma-Aldrich, MO, USA), an inhibitor of OAT transport [38]. The cell cultures were washed 3 times with ice-cold HBSS and were disrupted by incubating 100 μl 0.1 M NaOH for 10 min at 37°C. Fluorescence intensity was measured using GloMax® Explorer Multimode Microplate Reader (Promega, Madison, WI, USA), and expressed in arbitrary units (a.u.).

### 2.7 Bioengineering tubuloid-derived kidney tubules

For kidney tubule engineering, microPES (polyethersulfone) hollow fiber membranes were used, sterilized with 70% (v/v) EtOH incubation for 30 min as described previously [16,39,40]. An initial coating was performed by incubating the fibers with 10 mM L-3,4-dihydroxyphenylalanine (L-DOPA) (Sigma-Aldrich, MO, USA) dissolved in 10 mM Tris buffer (pH 8.5) at 37 °C for 5 h [39]. The second layer of coating was given by fiber incubation with human collagen IV (C6745-1 ml, 25 μg.ml^−1^) for 1 hour at 37 °C.

The tubuloids were dissociated in single cells as described above and seeded on double-coated fibers (length 2 cm) using 1 × 10^6^ cells/fiber and incubated at 37 °C and 5% v/v CO_2_ for 16 h. After this, the unattached cells were removed and the attached cells were maintained in culture with expansion medium until covering the entire surface of the fiber. Posteriorly, the cell cultures on fibers were submitted to the differentiation protocol for the different experimental conditions (Figure 1a).

### 2.8 Immunofluorescent staining

Tubuloid-derived cell cultures in 6-well plates or fibers were fixated with paraformaldehyde 4% for 15 min. In the case of 3D conformation, the tubuloids were initially incubated with Dispase to remove the BME followed by fixation with paraformaldehyde 4%. The cells were permeabilized in 0.3% (v/v) triton X-100 in HBSS for 10 min and incubated with blocking solution containing 2% (w/v) bovine serum albumin fraction V (Roche, Woerden, The Netherlands) and 0.1% (v/v) tween-20 in HBSS for 30 min. The cells were incubated with primary antibodies diluted in block solution against OAT1 (ab135925, Abcam), the tight junction protein zonula occludens 1 (ZO-1; ab216880, Abcam), Na^+^/K^+^-ATPase (kindly provided by Dr JB Koenderink, Radboudumc, Nijmegen, the Netherlands [41]), acetylated α-tubulin (T6793, Sigma-Aldrich, MO, USA). The secondary antibodies used were the anti-rabbit-Alexa488 and the anti-mouse-Alexa-594 conjugates (Life Technologies Europe BV, Bleiswijk, The Netherlands). Phalloidin-iFluor 488 Reagent (ab176753) was used to evaluate F-actin expression. Nuclei were stained with ProLongTM Gold antifade reagent containing DAPI (Life Technologies Europe BV, Bleiswijk, The Netherlands). Images were obtained using confocal microscopy (Leica TCS SP8 X, Leica Biosystems, Amsterdam, The Netherlands) and the software Leica Application. To measure the cilia density, Multiple Z stacks images from immunofluorescent staining for acetylated α-tubulin were analyzed with ImageJ software to measure the total perimeter [40].

### 2.9 RNA isolation and quantitative real-time polymerase chain reaction (qRT-PCR)

mirVana RNA isolation kit (Thermo Fisher Scientific, MA, USA) was used for RNA extraction from tubuloid and EV-OAT1. RNA quantification was measured spectrophotometrically using NanoDrop™ OneC Spectrophotometer (Thermo Fisher Scientific, MA, USA) mRNA expression was assessed using a High-Capacity cDNA Reverse Transcription Kit (Applied Biosystems) and iQ SYBR Green Supermix (Bio-Rad, Hercules, CA, USA). Negative cDNA controls (no cDNA) were cycled in parallel with each run. qRT-PCR was done with a CFX96 Real-Time PCR Detection System (Bio-Rad). All the sequence-specific oligonucleotide primers were obtained from Biolegio (Nijmegen, The Netherlands). See Table S1 for primers sequences.

### 2.10 Western blotting

Protein extraction from tubuloid and EVs was performed using Ripa Buffer supplemented with Halt™ Protease and Phosphatase Inhibitor Cocktail (Thermo Fisher Scientific, MA, USA). The protein concentration was measured using Pierce™ BCA Protein Assay Kit and GloMax® Explorer Multimode Microplate Reader (Promega, Madison, WI, USA). Expression of OAT1, Na^+^/K^+^-ATPase and CD63 were measured by Western blotting. The OAT1 antibody (ab131087) was purchased from Abcam (Cambridge, UK), Na^+^/K^+^-ATPase (given by Dr JB Koenderink) [41] and CD63 (sc-5275) was purchase from Santa Cruz Biotechnology (Dallas, TX, USA). The β-actin antibody was used as a loading control (ab8226, Abcam). The secondary antibodies used were goat anti-rabbit and rat anti-rabbit IgG-HRP (Dako, Santa Clara, CA, USA). Proteins were detected by chemiluminescence using the Clarity™ Western ECL Substrate coupled to a ChemiDoc XRS+(Bio-Rad, Hercules, CA, USA). Quantification of Western blots relied on ImageJ software [42].

### 2.11 Statistical analysis

Statistical analyses were performed using Student t-test or one-way analysis of variance (ANOVA) test with Tukey’s post-test, were appropriate. Statistical significance was set at P < 0.05. Data were plotted and analyzed using GraphPad Prism 5.0 (GraphPad Software, San Diego, CA USA). All data are expressed as mean ± SEM.

## 3. RESULTS

### 3.1 Characterization of EVs and their uptake by the tubuloids

To establish the experimental conditions (Figure 1a), an initial characterization of EV-OAT1 was performed and it was found that mature ciPTEC-OAT1 secrete a heterogeneous population of EVs, ranging from 20 nm to 594 nm, with a mean value of 131 nm. A representative image of EV-OAT1 size distribution is presented in Figure 1b. Western blotting of the EVs showed the presence of OAT1 amongst its cargo (Figure 1c). Moreover, qRT-PCR analysis revealed the presence of mRNA of OAT1 (*SLC22A6*; 2^−ΔCt^: 3.27 ± 0.55, relative to the housekeeping gene HPRT1), indicating that OAT1 can be delivered into the tubuloids as mRNA and protein constructs. Proteomic analysis of EVs confirmed the presence of the exosome population by markers like CD9, CD63, CD81 and TSG101 [43] in the top 50 most expressed proteins in EVs in the EVpedia database (Table 1 and S2), of which CD63 expression was also confirmed by Western blotting (Figure 1d). Another important protein identified in the EVs is the Na^+^/K^+^-ATPase Transporting Subunit Alpha 1 (ATP1A), an enzyme encoded by the *ATP1A1* gene that is crucial for maintaining the electrochemical gradients of sodium and potassium ions across the plasma membrane. These gradients are essential for osmoregulation and for sodium-coupled transport of a variety of organic and inorganic molecules, including the transport of organic anions by OAT1 [44]. The abundance of Na^+^/K^+^-ATPase is also an indication of cell polarization [45]. Western blotting of the EV-OAT1 confirmed its presence (Figure 1e).

**Table 1.** Top 50 proteins most expressed in the EVs identified in the EVpedia database.

| Index | Gene | Protein name | Identified | Index | Gene | Protein name | Identified |
| --- | --- | --- | --- | --- | --- | --- | --- |
| 1 | PDCD6IP | Programmed cell death 6-interacting protein | 399 | 26 | MSN | Zinc finger protein MSN2 | 266 |
| 2 | GAPDH | Glyceraldehyde-3-phosphate dehydrogenase | 377 | 27 | ATP1A1 | Na(+)/K(+) ATPase alpha-1 subunit | 266 |
| 3 | HSPA8 | Heat shock cognate 71 kDa protein | 363 | 28 | PRDX1 | Peroxiredoxin-1 | 263 |
| 4 | ACTB | Actin, cytoplasmic 1 | 350 | 29 | MYH9 | Myosin-9 | 262 |
| 5 | ANXA2 | Annexin A2 | 337 | 30 | EZR | Ezrin | 262 |
| 6 | CD9 | CD9 antigen | 328 | 31 | CD81 | CD81 antigen | 262 |
| 7 | PKM | Pyruvate kinase muscle isozyme | 327 | 32 | ANXA6 | Annexin A6 | 260 |
| 8 | HSP90AA1 | Heat shock protein HSP 90-alpha | 327 | 33 | FLOT1 | Flotillin-1 | 259 |
| 9 | ENO1 | Alpha-enolase | 327 | 34 | YWHAB | 14-3-3 protein beta/alpha | 258 |
| 10 | ANXA5 | Annexin A5 | 313 | 35 | LDHB | L-lactate dehydrogenase B chain | 258 |
| 11 | HSP90AB1 | Heat shock protein HSP 90-beta | 306 | 36 | SLC3A2 | 4F2 cell-surface antigen heavy chain | 257 |
| 12 | CD63 | CD63 antigen | 306 | 37 | GNB1 | Guanine nucleotide-binding protein G(I)/G(S)/G(T) subunit beta-1 | 257 |
| 13 | YWHAZ | 14-3-3 protein zeta/delta | 301 | 38 | PFN1 | Profilin-1 | 256 |
| 14 | YWHAE | 14-3-3 protein epsilon | 300 | 39 | TSG101 | Tumor susceptibility gene 101 protein | 255 |
| 15 | EEF1A1 | Elongation factor 1-alpha 1 | 295 | 40 | YWHAQ | 14-3-3 protein theta | 254 |
| 16 | PGK1 | Phosphoglycerate kinase 1 | 291 | 41 | GNAI2 | Guanine nucleotide-binding protein G(i) subunit alpha-2 | 252 |
| 17 | CLTC | Clathrin heavy chain 1 | 283 | 42 | CLIC1 | Chloride intracellular channel protein 1 | 251 |
| 18 | PPIA | Peptidyl-prolyl cis-trans isomerase A | 278 | 43 | ANXA1 | Annexin A1 | 251 |
| 19 | SDCBP | Syntenin-1 | 277 | 44 | ITGB1 | Integrin beta-1 | 250 |
| 20 | ALDOA | Fructose-bisphosphate aldolase A | 275 | 45 | LDHA | L-lactate dehydrogenase A chain | 249 |
| 21 | EEF2 | Protein-lysine N-methyltransferase EEF2KMT | 274 | 46 | FASN | Type I Fatty Acid Synthase | 248 |
| 22 | ALB | Albumin | 274 | 47 | CDC42 | CDC42 small effector | 248 |
| 23 | TPI1 | Triosephosphate isomerase | 270 | 48 | RAP1B | Ras-related protein Rap-1b | 242 |
| 24 | VCP | Transitional endoplasmic reticulum ATPase | 269 | 49 | CCT2 | T-complex protein 1 subunit beta | 242 |
| 25 | CFL1 | Cofilin-1 | 268 | 50 | YWHAG | 14-3-3 protein gamma | 240 |

To determine whether the EVs could be internalized by the tubuloids, EVs isolated from Vybrant DiI-stained ciPTEC-OAT1 (Figure 1f) were used, which showed that within 24 h of incubation the tubuloids internalized EV-OAT1 (Figure 1g) and widely distributed in the tubuloids (Figure 1h).

### 3.2 CM-OAT1 and EV-OAT1 enhance the expression of transepithelial transporters in tubuloids

Evaluation of OAT1 expression in tubuloids revealed that CM-OAT1 induced OAT1 mRNA levels by 3.3-fold and protein levels by 4.2-fold, compared to control (Figure 2a). Interestingly, the tubuloids did not present an increase in OAT1 mRNA when incubated with CM-OAT1 depleted of EVs confirming that the EVs and their cargo are responsible for the upregulation by CM-OAT1 (Figure 2a). Such enhancement may be given either by a regulatory action of factors within EVs that stimulate OAT1 expression or by a direct transfer of mRNA and or protein. To better understand the mechanism involved in OAT1 upregulation, we exposed tubuloids to two different concentrations of EV-OAT1: 1 stimulation (5×10^8^ particles/well; TUB EV-OAT1 1st) or 3 stimulations in a single day (1.5×10^9^ particles/well; TUB EV-OAT1 3st), in the first 24 h of tubuloids differentiation phase (Figure 2b). The tubuloids stimulated with a single dose or three doses showed, respectively, a 1.8- and 3-fold increase at the OAT1 mRNA levels, indicating a dose-depended response related to the delivery of EV-OAT1 cargo.

**Figure 2.**
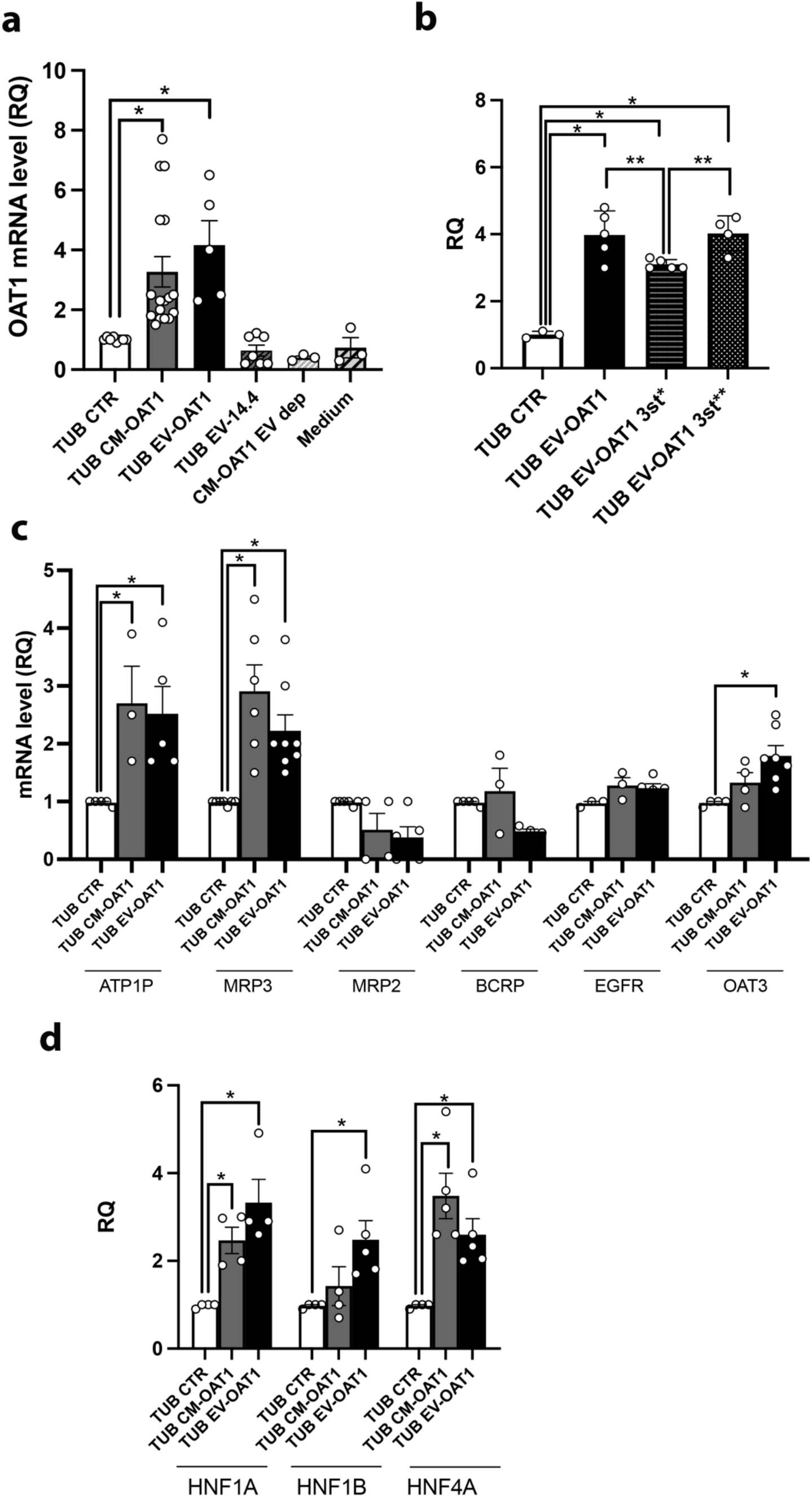
EV-OAT1 promote changes in epithelial transporters and transcription factors genes in tubuloids. (a) Conditioned medium and extracellular vesicles from ciPTEC-OAT1 (CM-OAT1 and EV-OAT1) upregulate OAT1 in tubuloids (TUB). The graph shows the changes in the mRNA levels in the tubuloids in different experimental conditions indicated in the abscissa (CTR represents standard differentiation protocol; EV-14.4 indicates EVs derived from ciPTEC 14.4; EV dep indicated CM-OAT1 depleted of EVs; Medium indicates the differentiation medium submitted to the same process of concentration for EVs and that was incubated with the tubuloids). (b) EV-OAT1 promote OAT1 upregulation in the tubuloids in a dose-dependent manner. The graph indicates the mRNA OAT1 levels stimulates with EV-OAT1 with a single dose (TUB EV-OAT1 1st) or three doses in a single stimulation (TUB EV-OAT1 3st). (c) The graph shows the changes in OAT1 mRNA levels in the tubuloids after different stimulation timepoints. The abscissa indicates the condition: TUB EV-OAT1 indicates normal stimulation protocol with administration of 3 doses in 3 different days within 7 days; 3st* indicates the stimulation of 3 doses in a singles stimulus in the beginning of differentiation protocol (day 1), while 3st** indicates the same single stimulus at the end of the protocol (day5). (d) CM-OAT1 and EV-OAT1 promote the upregulation of other epithelial transporters in tubuloids. The graph shows the mRNA levels of epithelial transporters (ATP1, MRP3, MRP2, BCRP and OAT3) of tubuloids cultured in the different experimental conditions (abscissa). (e) Transcription factors associated with drug transport are upregulated by CM-OAT1 and EV-OAT1. The graph shows the mRNA levels of transcription factors in the tubuloids (HNF1A, HNF4A, HNF1B). In all graphs, the data is expressed in relative quantification (RQ) with respect to the control condition (TUB CTR) (n = 5). Data represent mean ± SEM, (*) p<0.05 with respect to TUB CTR group and (**) p<0.05 with respect to TUB EV-OAT1 3st* group.

Furthermore, we evaluated if such modulatory effect in tubuloids could be maintained for a long period after EV-OAT1 stimulation (Figure 2c). When the 3 stimulations (1.5×10^9^ particles/well) are given in day 1 of the maturation phase and tubuloids are maintained in culture without further stimulation for 7 days (TUB EV-OAT1 3st*), the levels of OAT1 are still higher compared to control condition (3-fold increase). However, such levels remained lower than the 3 stimuli along 7 days (TUB EV-OAT1), even though the total amount of EV-OAT1 administrated was the same. Differently, when the 3 EV-OAT1 stimulations are added in the end of differentiation protocol (TUB EV-OAT1 3st**), day 5, the OAT1 levels are similar to the 3 stimuli along 7 days (4-fold increase). These results indicate that OAT1 upregulation mediated by EVs is not exclusively given by direct transfer, once the expression levels are kept higher even 7 days after EV administration. In addition, the stimulation performed closer to the end of differentiation protocol may be more effective to induce such phenotype. It is worth to mention that despite the capacity of EV-OAT1 to regulate tubuloids OAT1 expression, the transfer of OAT1 mRNA cargo is not excluded. The amount of OAT1 mRNA present inside the EVs is compatible with the increased level observed in the tubuloids after incubation with EV-OAT1 (Figure S1).

Since ciPTEC-OAT1 overexpress OAT1, we evaluated the effects on tubuloids of EVs derived from the parent cells that do not express OAT1 (ciPTEC-14.4 [31]). Size distribution analysis of EVs derived from ciPTEC-14.4 (EV-14.4) showed a similar profile as obtained for EV-OAT1 (ranging from 22 nm to 575 nm, with a mean value of 180 nm), but lack the capability of inducing OAT1 expression in tubuloids (Figure 2a), indicating that the high expression of the transporter in ciPTEC-OAT1 is a vital element for the modulatory effect of EVs. Moreover, EV-OAT1 isolated with additional purification step using ExoQuick-TC also promoted the expression of OAT1 in the tubuloids (Figure S2), confirming that the effects are exclusively mediated by EV-OAT1 and that EVs act as transfection agents in cargo delivery.

To explore further the changes promoted by EV-OAT1, we analyzed the mRNA levels of other transporters linked to OAT1 and associated with the excretion of metabolic waste and drug handling, including *ATP1A, ABCC2* (MRP2) and *ABCC3* (MRP3), *ABCG2* (BCRP) and *SLC22A8* (OAT3; Figure 2d). Of these transporters, *ATP1A* and *ABCC3* were equally increased by CM-OAT1 and EV-OAT1 when compared to control (*ATP1A*; CM: 2.7-fold increase; EV: 2.5-fold increase; *ABCC3*; CM: 2.9-fold increase; EV: 2.2-fold increase), whereas the expression levels for *ABCC2* and *ABCG2* were not altered. Interestingly, exposure of tubuloids to EV-OAT1 resulted in a 1.8-fold increase in OAT3 mRNA levels, which was not observed in CM-OAT1 exposures. However, the increase in OAT3 was not as pronounced as observed for OAT1, indicating once more that the transporter expression levels in the originator cells of EVs is key in the modulation of the tubuloids as EV recipients.

### 3.3 CM-OAT1 and EV-OAT1 modulate the expression of transcription factors associated with the regulation of epithelial transporters and kidney tubular maturation

To better understand the regulatory action of CM and EVs derived from ciPTEC-OAT1, we analyzed the expression of some transcriptional factors known to regulate tubule epithelial cell maturation and the expression of epithelial transporters [46] (Figure 2e). The mRNA levels of the transcriptional factors hepatocyte nuclear factor 1 alpha, 4 alpha and 1 beta (*HNF1A, HNF4A* and *HNF1B*) revealed that CM-OAT1 and EV-OAT1 were capable of regulating these genes in the tubuloids. CM-OAT1 promoted the expression of *HNF1A* (2.5-fold) and *HNF4A* (3.5-fold), but not *HNF1B*. In contrast, isolated EV-OAT1 led to the upregulation of all three transcription factors in the tubuloids compared to control (*HNF1A*: 3.3-fold; *HNF4A:* 2.6-fold; *HNF1B:* 2.5-fold).

### 3.4 CM-OAT1 and EV-OAT1 improve tubuloid OAT1 protein expression, localization and transport capacity

The localization of OAT1 in the basolateral membrane is crucial for the vectorial transport of substrates such as metabolic wastes, but it is also an indication of epithelial cell polarization. Immunostaining of tubuloids after exposure to CM-OAT1 and EV-OAT1 demonstrated an increased polarized expression of OAT1 in the basolateral membrane of the cells, facing the outer part of the tubuloids (Figure 3a). The tubuloids cultured under standard differentiation condition presented a more disperse localization of OAT1, although some basolateral foci of OAT1 can be observed as well. Quantification by Western blotting revealed that CM-OAT1 and EV-OAT1 induced an increase in OAT1 protein (Figure 3b), but also in Na^+^/K^+^-ATPase (Figure 3c), confirming a polarized maturation of the tubuloids.

**Figure 3.**
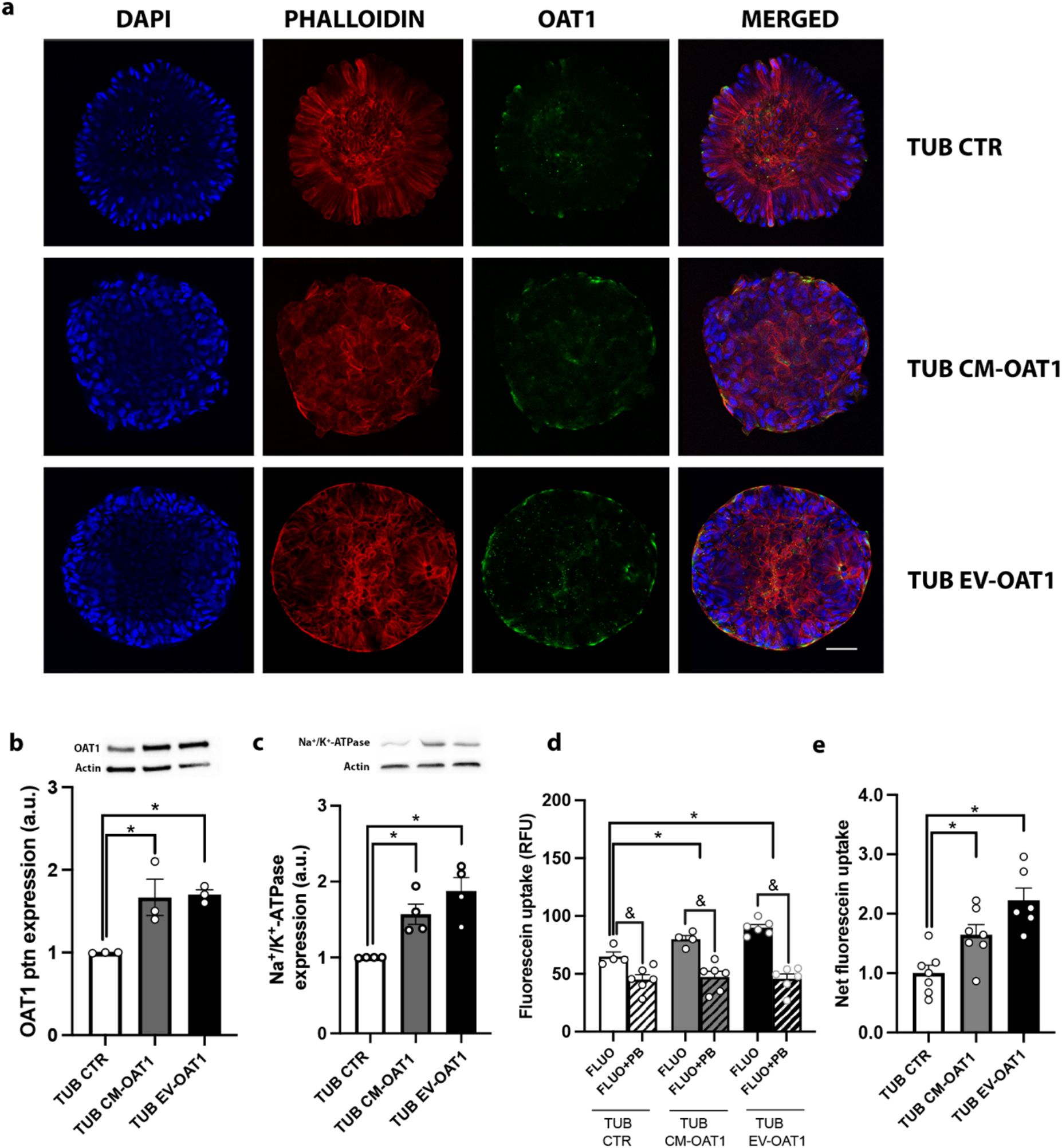
EV-OAT1 and CM-OAT1 support functional maturation of tubuloids. (a) Representative confocal images of OAT1 localization in the tubuloids under different experimental conditions. The nuclei of the tubuloid cell were stained with DAPI (in blue). The tubuloid spatial organization is indicated by actin disposition, stained with phalloidin (in red). OAT1 localization is observed by the green staining. The last column represents the merge of the three images of each respective experimental condition (scale bar = 50 μm). (b) OAT1 protein expression in the tubuloids. Upper panel shows representative images of Western blot for OAT1 and Actin. The graph shows the quantification of OAT1 expression in the Western blotting (n = 3). (c) Na^+^/K^+^-ATPase protein expression in the tubuloids. Upper panel shows representative images of Western blot for Na^+^/K^+^-ATPase and Actin. The graph shows the quantification of Na^+^/K^+^-ATPase expression after Western blotting (n = 3). (d) Fluorescein uptake capacity by the tubuloids. The graph shows the intracellular fluorescence intensity of tubuloids after 10 min incubation with fluorescein (FLUO), in the presence or absence of probenecid (PB) (OATs inhibitor) (n = 4). The fluorescence intensity is expressed as arbitrary units (a.u.). (e) Net fluorescent uptake specific to OAT in tubuloids. The graph shows the increase of fluorescein uptake of tubuloids cultured with CM-OAT1 or EV-OAT1 in respect to tubuloids under standard differentiation condition. The specificity of transport was given by the difference in the fluorescence intensity between the presence and absence of probenecid. The data is presented as ration in respect to TUB CTR condition (n = 4). In all graphs, data represent mean ± SEM, (*) p<0.05 compared to TUB CTR group, (&) p<0.05 compared to FLUO+PB for each experimental condition.

Increased expression of OAT1 at the basolateral membrane should ultimately lead to an increase in the functional capacity of the tubuloids to transport organic anions. To this end, we evaluated the transport efficiency of OAT1 through fluorescein uptake by the tubuloid cells (Figures 3d, e) and demonstrated an increase in the intracellular fluorescence intensity when compared to control (Figure 3d), which was sensitive to probenecid, a known OAT1 inhibitor, confirming active transporter-mediated uptake. This was more pronounced when tubuloids were matured in the presence of EV-OAT1, supporting EVs’ potential to functionally mature tubuloids (Figure 3e). It is worth to mention that OAT3 may also facilitate probenecid-sensitive fluorescein transport, but with a lower affinity [18]. The increased OAT1 expression in tubuloids cultured with CM-OAT1 together with the absence of changes in the OAT3 mRNA levels (Figure 2b), however, argue for the improvement in transport capacity by OAT1.

### 3.5 Cell maturation processes drive maturation of tubuloids by EVs

To better understand the mechanism through which EVs modulate tubuloids, we performed a proteomic analysis and compared the profiles of the cargo of EV-OAT1 with those of EV-14.4 (Figure 4a, Table S2). We identified 964 proteins upregulated or exclusively expressed in EV-OAT1, 367 proteins upregulated or exclusively present in EV-14.4 (also defined as downregulated in EV-OAT1) and 597 proteins commonly expressed in both EVs (Figure 4b). Functional enrichment analysis of the genes associated with the 964 upregulated proteins in EV-OAT1 indicated that the identified proteins are mainly associated with protein metabolism, cellular metabolism and energy pathways (Figure 4c). Moreover, the biological pathways associated with the upregulated proteins showed a regulatory capacity of EVs to modulate gene expression (e.g., mRNA splicing, translation initiation and termination (Figure 4d)).

**Figure 4.**
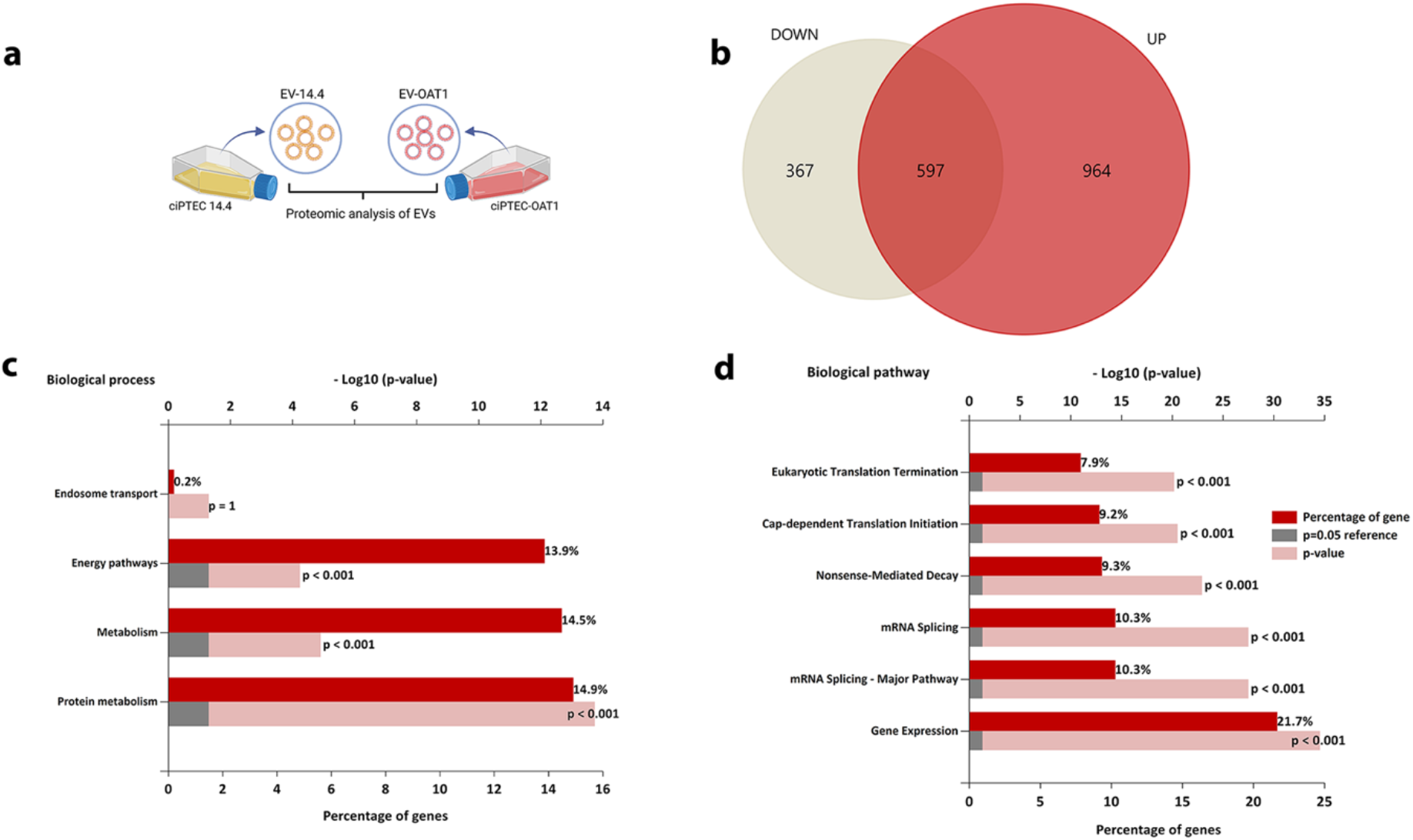
Proteomic analysis of EV-OAT1. (a) Representative scheme of the comparison of proteins presents in EV-OAT1 and EV-14.4. (b) The Venn diagram shows the proteins that are exclusively present or upregulated in EV-OAT1 (UP), commonly expressed and absent or downregulated in EV-OAT1 (DOWN) compared to EV-14.4. (c) The graph indicates the biological processes associated with the exclusively present or upregulated proteins in EV-OAT1. (d) The graph indicates the biological pathways associated with the exclusively present or upregulated proteins in EV-OAT1. The abscissa indicates −Log10(p-value).

Furthermore, the proteome of tubuloids cultured under the three different experimental conditions (CTR, CM-OAT1 and EV-OAT1) was compared (Figure 5, Tables S3-S5). The data obtained showed that 188 proteins were upregulated in the tubuloids cultured with CM-OAT1 and 245 proteins were upregulated in tubuloids cultured with EV-OAT1, compared to control (Figure 5a, b). Among the identified proteins, 118 were shown to be commonly upregulated in tubuloids cultured with CM-OAT1 and EV-OAT1, including neutrophil gelatinase-associated lipocalin (Lcn2/NGAL) and Plexin B2 (Plxnb2) known regulators of kidney maturation [47,48]. Functional enrichment analysis of the genes associated with these proteins revealed that they are mainly associated with energy pathway and cellular metabolism (Figure 5c), similar to those observed in the EVs proteomic analysis. Notably, the mesenchymal-to-epithelial transition (MET) is one of the key pathways triggered by EVs (Figure 5d).

**Figure 5.**
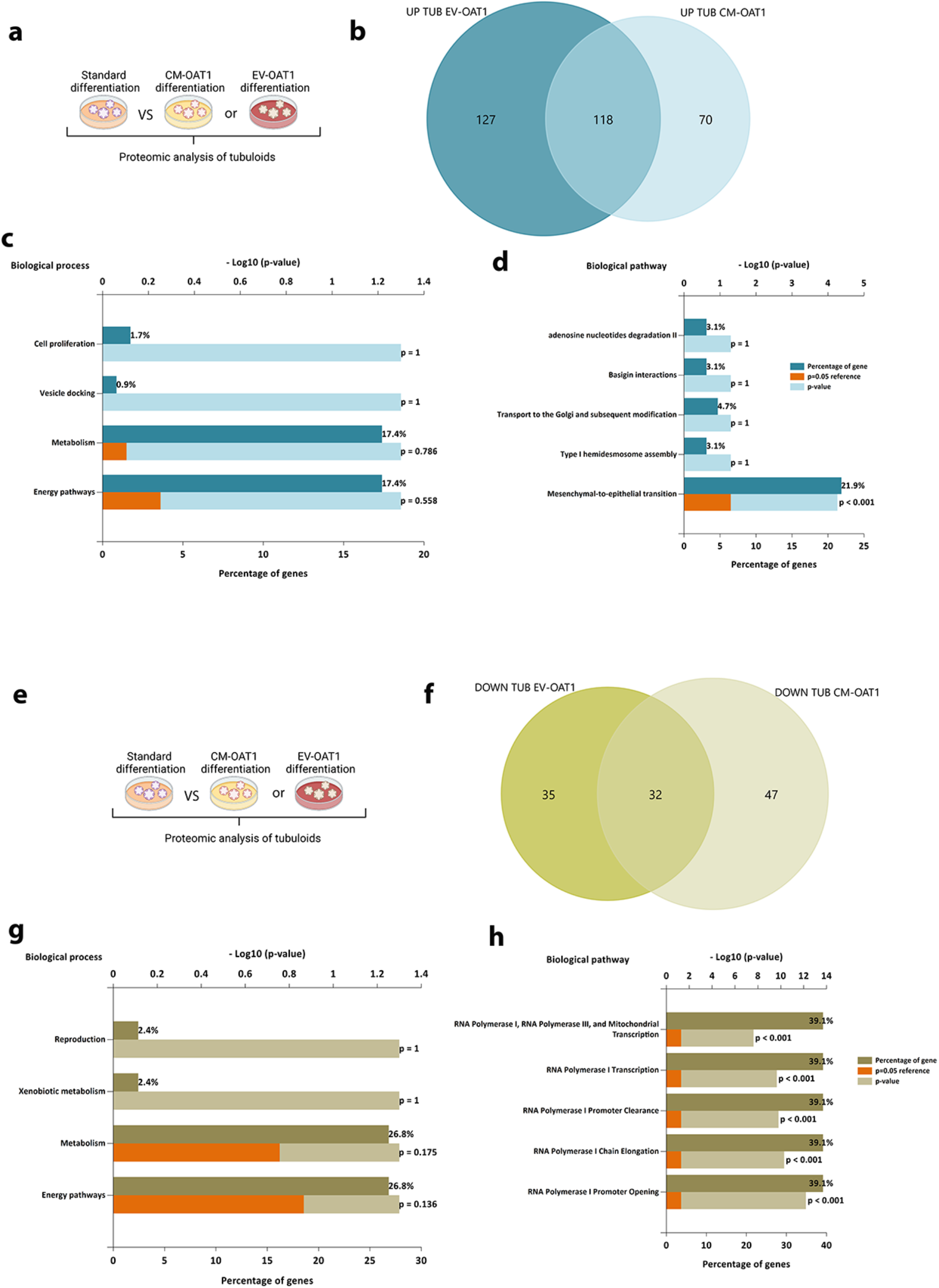
Pathways involved in the tubuloid changes promoted by CM-OAT1 and EV-OAT1. (a) Representative scheme of the comparison of proteins exclusively upregulated in tubuloids cultured with CM-OAT1 or EV-OAT1. (b) The Venn diagram shows the upregulated proteins in the tubuloids incubated with CM-OAT1 or EV-OAT1 as compared to tubuloids cultured under standard differentiation protocol. (c) The graph indicates the biological processes associated with the commonly upregulated proteins in tubuloids cultured with CM-OAT1 or EV-OAT1. (d) The graph indicates the biological pathways associated with the commonly upregulated proteins in tubuloids cultured with CM-OAT1 or EV-OAT1. The abscissa indicates −Log10(p-value). (e) Representative scheme of the comparison of proteins exclusively downregulated in tubuloids cultured with CM-OAT1 or EV-OAT1. (f) The Venn diagram shows the downregulated proteins in the tubuloids incubated with CM-OAT1 or EV-OAT1 compared to tubuloids cultured under standard differentiation protocol. (g) The graph indicates the biological processes associated with the commonly downregulated proteins in tubuloids cultured with CM-OAT1 or EV-OAT1. (h) The graph indicates the biological pathways associated with the commonly downregulated proteins in tubuloids cultured with CM-OAT1 or EV-OAT1. The abscissa indicates −Log10(p-value).

Furthermore, 79 proteins were downregulated in tubuloids cultured with CM-OAT1 and 67 proteins in the presence of EV-OAT1 (Figures 5e, f). The common 32 downregulated proteins were mainly associated with cellular metabolism and energy supply, although not statistically significant (Figure 5g). The biological pathways associated with the reduced proteins indicate a strong relation with the regulation of RNA transcription (e.g., RNA polymerase I promoter opening, chain elongation and transcription) (Figure 5h). Among the downregulated proteins, the histone H3 subunits were mainly reduced by EVs, indicating a role in regulation of chromatin organization and DNA replication (Table S4) [49].

### 3.6 EV-treated tubuloids form polarized kidney tubules on hollow fiber scaffolds

One of the potential applications of tubuloids in regenerative medicine is the creation of functional units in a bioartificial kidney to replace, aid or enhance kidney function in affected patients. We previously established a physiological model of a polarized kidney epithelial cell layer by seeding ciPTECs onto a hollow-fiber scaffold that allows transepithelial transport under perfusion [16,40]. Here, we used a similar approach for the tubuloid-derived cells to bioengineer kidney tubules (Figure 6a). Tubuloid cells adhered to biofunctionalized hollow fibers and proliferated, thereby covering the entire fiber surface. Next, the tubules were cultured under one of the differentiation protocols to establish a tight 3-dimensional monolayer, which was confirmed by immunofluorescent staining of the tight-junction protein ZO-1, with no differences among the three experimental conditions (Figure 6b). Furthermore, localization of Na^+^/K^+^-ATPase in the basolateral membrane and cilia structure formation at the apical region confirmed adequate polarization of the tubuloid-derived monolayers (Figure 6c – f). Again, culturing the kidney tubules in the presence of CM-OAT1 or EV-OAT1 enhanced the cell polarity process as indicated by an increased cilia density determined by the ratio of the cilia total perimeter and the number of cells (Figure 6g).

**Figure 6.**
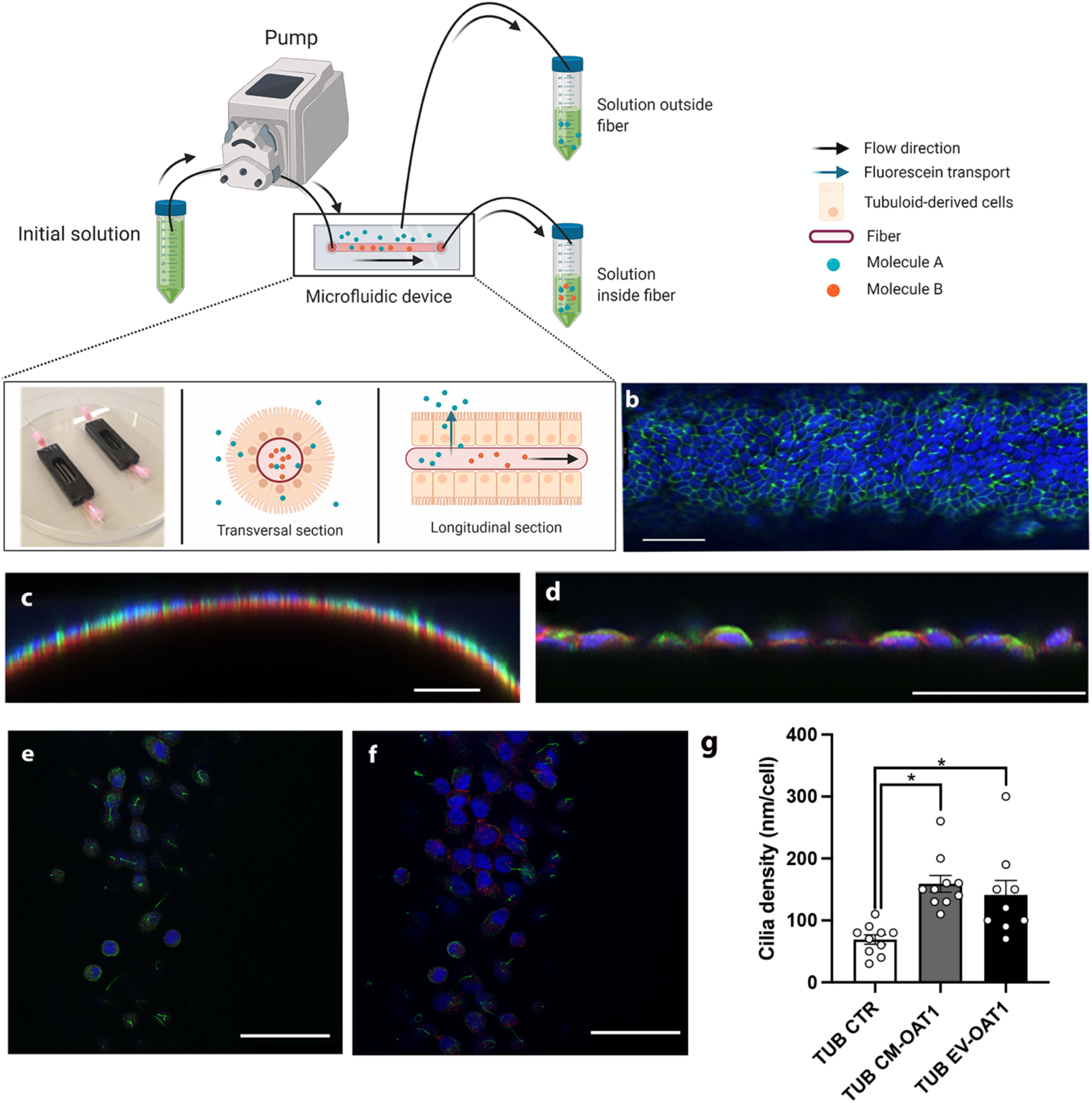
Tubuloid-derived cell form tight monolayer on microfluidic hollow fiber-based platform and CM-OAT1 and EV-OAT1 improve cilia density. (a) Scheme of the microfluidic hollow fiber-based platform and how tubuloid-derived cells are organized, and selectively transport molecules from one compartment to another. (b) Representative confocal image of tubuloids-derived cells cultured on hollow fibers in the presence of EV-OAT1. The ZO-1 immunostaining (in green) shows the presence of a tight epithelial monolayer on the fiber. (c) Representative y-z confocal image of the curved surface of the fiber and the polarized tubuloid-derived cells cultured with EV-OAT1. The acetyl-α-tubulin immunostaining in the apical region (in green) indicates the cilia structure formation and Na^+^/K^+^-ATPase is localized to the basolateral region (in red). (d) Higher magnification of tubuloids cells on the fibers with α-tubulin (in green) and Na^+^/K^+^-ATPase (in red) staining. (e) The cilia structures (in green) are mainly observed when the apical region is in focus, while in (f) the presence of Na^+^/K^+^-ATPase (in red) is mainly observed in the basolateral region. In all images, scale bar = 50 μm. (g) Quantification of total cilia perimeter present in the tubuloids-derived cells under the different experimental conditions. The graph shows the total cilia perimeter with respect to the total number of cells. Data represent mean ± SEM, (*) p<0.05 compared to TUB CTR group.

## 4. Discussion

EVs have been described as an essential element in intranephron communication and kidney regeneration, mediated by the transfer molecules and from one cell type to another [50]. In this study, we propose a novel role for EVs as molecular messengers for functional maturation of kidney tubular organoids *in vitro*. We showed that EVs derived from mature kidney cells overexpressing OAT1 (EV-OAT1) improved the tubuloids phenotype by triggering various biological pathways associated with epithelial maturation. As a result, tubuloids showed enhanced polarization, increased formation of cilia structures, and improved epithelial transport capacity with increased mRNA levels of several proximal tubular transporters, including OAT1. The evolutionarily well conserved solute carrier coordinates the active secretion of a broad range of endogenous and exogenous substances. Unfortunately, the transporter is rapidly lost in primary kidney cell cultures and not functionally present in tubuloids. The use of EVs provide a tool to induce the transporter expression, which amplifies the tubuloids applications. Concomitantly, *HNF1A, HNF4A* and *HNF1B* were upregulated, three master regulatory transcription factors towards kidney lineage differentiation [51], kidney tubular epithelial maturation [52] and drug transporters, including OAT1 and OAT3 [50, 53–55]. It should be noted, however, that OAT1 upregulation in tubuloids was much more pronounced upon CM and EV exposure than OAT3, which is likely due to the high expression levels of the transporter in ciPTEC-OAT1, whereas OAT3 is absent in this cell line. Western blotting and qRT-PCR analysis showed that EV-OAT1 cargo contains OAT1 as both mRNA and as protein. Therefore, a direct transfer of OAT1 cargo most likely drives the upregulation of OAT1, although indirect transcriptional regulation should not be excluded as OAT3 was also upregulated in the tubuloids in the presence of EV-OAT1.

The observed upregulation of Na^+^/K^+^-ATPase is of great importance for functional OAT1 as organic anion transport is tertiary coupled to the sodium gradient generated by this pump [56]. Again, it remains to be elucidated whether the joint upregulation of OAT1 and Na^+^/K^+^-ATPase is mediated indirectly by induced regulatory pathways or through a direct transfer of mRNA and protein, or both.

Besides proximal tubule cells, tubuloids also contain cells from the distal tubule, loop of Henle, and collecting duct epithelium, which take up EVs [10]. Interestingly, EV exposure increased the expression of the basolateral transporter MRP3, an efflux pump expressed in kidney distal tubule cells [55], indicating that EV-OAT1 also promoted changes in cell types other than proximal tubule cells. This argues for EVs mediating cell responses not only via direct cargo transfer as MRP3 is not expressed in the originator cells.

To gain more insights into the effects of EVs on tubuloids, we performed proteomic analyses. The main biological processes modulated upon CM and EV exposures were energy pathways and cellular metabolism, similar to processes found in EV’s cargo analysis, which supports the relation between EV cargo composition and the response triggered in the target cells. Furthermore, pathway analysis indicated that mesenchymal-to-epithelial transition is one of the main pathways activated, which supports the phenotypic improvements observed (e.g., cellular polarization and increased cilia density).

Among the 964 upregulated proteins in tubuloids incubated with EVs, we identified neutrophil gelatinase-associated lipocalin (Lcn2/NGAL) that has been described to facilitate iron delivery into cells. Lcn2 regulates iron-sensitive genes that participate in mesenchymal-to-epithelial transition during the development of the proximal parts of the mammalian nephron [47,57]. Additionally, Plexin B2 (*Plxnb2*), a semaphorin receptor expressed in the pretubular aggregates and the ureteric epithelium in the developing kidney, was found upregulated [58]. Plxnb2-deficient mice were shown to present intrinsic defects of the ureteric epithelium, leading to reduced branching and proliferation. Moreover, this receptor is directly associated with the formation and organization of the collecting duct, a nephron segment that is also present in the tubuloids [10,48]. One of the most downregulated proteins upon EV exposure was histone H3. Histone H3 synthesis is coupled to DNA replication, providing material for the bulk of nucleosome assembly for the duplicated genome [59]. Thus, downregulation of histone H3 indicates a reduction in tubuloid cell proliferation. This is supported by the reduced expression of the proliferating cell nuclear antigen (PCNA), which is rapidly downregulated as kidney epithelial cells enter terminal differentiation and acquire functional characteristics during kidney development [60].

EVs have been vastly investigated for kidney disease treatment as regulators of several processes, including counteracting inflammatory responses, as anti-oxidative stress response, and prevention of cell death [32,33,61–63]. In this study, we demonstrated a new application potential for EVs in regenerative nephrology by supporting tubuloid maturation on a hollow fiber-based microfluidics device to be used as functional unit of a bioartificial kidney. Previously, such device has been shown to successfully recreate aspects of kidney tubular function by establishing a polarized monolayer capable of promoting active waste transport using ciPTEC-OAT1 [39,64]. The immortal nature of these cells, harboring two oncogenes, however, hamper their future clinical application in kidney replacement therapies. This limitation can be overcome by the use of tubuloids if their maturation can be supported. The present results show that tubuloids can be matured by EVs and used to form a tight epithelial monolayer on hollowfiber membranes with enhanced polarization given by increased cilia structure density. Cilia are sensory organelles located at the apical membrane that sense fluid flow and initiate calcium-based signaling to regulate tubular function and maintain an epithelial phenotype [65,66]. Moreover, cilia structures have been shown to play a critical role in the Hedgehog and Wnt signalling pathways that are also associated with epithelial development [67,68]. These findings indicate that EVs can be a key tool for the maturation of tubuloids and kidney tubule engineering. By bioengineering EVs with a finetuned cargo [61,69], tubuloids might be phenotypically modulated even further in the future to approach a near-physiological phenotype as found in native tissue. In this manner, the EVs may represent a key element in the development of cell-based bioartificial kidneys from tubuloids/organoids, guiding the differentiation and maturation process of different kidney cell types.

In conclusion, EVs from renal tubular epithelial cells are an important element in the functional maturation of tubuloids. The regulatory actions of EVs are associated with a mesenchymal-to-epithelial transition process that led to increased expression of various epithelial transporters, improved cell polarity and transport capacity of tubuloids. Therefore, EVs can be considered an additional requisite to support the kidney microenvironment *in vitro*, allowing the development of kidney models closer to human physiology. Such innovative approach advances preclinical models’ development and supports applications in disease modelling and drug screening as well as in kidney replacement therapies. Moreover, the use of EVs can be extended to other tissue models to be applied as a strategy to support the maturation of organoids derived from adult stem cells and iPSCs. Finally, since EV composition is directly associated with their regulatory effects, we believe that identification of regulatory molecules to bioengineer EVs and used as a delivery system can promote a refined regulation of organoid fate in a patient-specific manner.

## ACKNOWLEDGEMENTS

This work was supported by the partners of Regenerative Medicine Crossing Borders (Regmed XB). Powered by Health~Holland, Top Sector Life Sciences and Health; the Dutch Ministry of Education, Culture and Science; the Brazilian National Research Council (grant number 421916/2016-8); and the Carlos Filho Rio de Janeiro State Research Foundation (E-26/010.000981).

## CONFLICTS OF INTEREST

The authors report no conflicts of interest.

---

Figure S1. OAT1 mRNA levels within EV-OAT1 is compatible with the increased levels in tubuloids. The graph shows the total OAT1 mRNA levels present in EV-OAT1 and tubuloids that were previously isolated separately and then pooled to perform the qRT-PCR (TUB+EV-OAT1 MIX). The expression levels were compared to unstimulated tubuloids (TUB CTR). The data is expressed in relative quantification (RQ) in respect to the control condition (TUB CTR) (n = 3). Data represent mean ± SEM, (*) p<0.05 with respect to TUB CTR group.

Figure S2. EV-OAT1 additionally purified from the CM maintained the upregulation of OAT1 in tubuloids. The graph shows the changes in the mRNA levels in the tubuloids culture under standard differentiation protocol (TUB CTR) and in the presence EV-OAT1 that were further isolated from the remaining medium using ExoQuick-TC. The data is expressed in relative quantification (RQ) in respect to the control condition (TUB CTR) (n = 3). Data represent mean ± SEM, (*) p<0.05 with respect to TUB CTR group.

Table S1. List of primers. (Data can be provide under request)

Table S2. List of all proteins identified in EV-OAT1 and its expression levels in respect to EV-14.4. (Data can be provide under request)

Table S3. List of all proteins identified in TUB CM-OAT1 and its expression levels in respect to TUB CTR. (Data can be provide under request)

Table S4. List of all proteins identified in TUB EV-OAT1 and its expression levels in respect to TUB CTR. (Data can be provide under request)

Table S5. List of all proteins identified in TUB EV-OAT1 and its expression levels in respect to TUB CM-OAT1. (Data can be provide under request)

## Notes

### Competing Interest Statement

The authors have declared no competing interest.

